# Putative MFS transporter Rv1250 of *Mycobacterium tuberculosis* is involved in multidrug efflux activity

**DOI:** 10.1101/2025.08.06.668847

**Authors:** Debasmita Chatterjee, Ajit Ramesh Sawant, Aditya Prasad Panda, Sadhana Roy, Somdeb Bose DasGupta, Anindya Sundar Ghosh

## Abstract

Drug-resistant *Mycobacterium tuberculosis* is one of the leading causes of global mortality. Mechanisms, such as slow uptake of drugs along with cell wall impermeability and active efflux, are some of the concerning reasons leading to drug resistance. Efflux pumps actively transport a wide variety of drugs and toxins away from the target site, which is considered an emerging cause for the failure of anti-tubercular medications and treatment. In this study, we report that the ability of Rv1250, a probable MFS-type transporter, influences the extrusion of multiple structurally unrelated classes of drugs, enhances the biofilm formation in E. coli and *Mycobacterium smegmatis*, and facilitates the survival of *M. smegmatis* cells inside the macrophage during antibiotic stress. Interestingly, *in trans*, the expression of rv1250 decreased the susceptibility of host cells to several structurally unrelated antibiotics, ranging from fluoroquinolones to aminoglycosides, beta-lactams, and anti-tubercular drugs, thus indicating its involvement in imparting intrinsic drug tolerance. In addition, the increased efflux of EtBr, norfloxacin, and Bocillin FL from host cells expressing rv1250 was revealed by the host’s ability to confer a lower level of antibiotic accumulation. Moreover, the expression of rv1250 resulted in the enhancement of biofilm formation. Overall, we conclude that Rv1250 of *Mycobacterium tuberculosis* might facilitate the survival of host cells under antimicrobial stress.

**One sentence summary:** Role of Rv1250 as an efflux pump

## Introduction

Tuberculosis (TB) is one of the leading causes of death from a single infectious agent worldwide. On record, more than 10 million people continue to fall ill due to TB ^1^. The occurrence of multidrug-resistant TB (MDR-TB), extremely drug-resistant TB (XDR-TB) are of grave concern^2^. *M. tuberculosis* (*Mtb*) survives inside the host by evading the host defence system and by resisting the arsenal of antimicrobial therapies. Along with cell wall impermeability, chromosomal mutations, the presence of antibiotic-hydrolysing enzymes, and active efflux pumps can also contribute to the drug tolerance mechanism of *M. tuberculosis* ^3,4^. The efflux pumps are transmembrane proteins responsible for the reduction of drug-target interaction by the export of drugs, metabolites and toxic substances out of the cells ^5,6^. The pumps are categorised into two classes based on their energy sources. The primary active transporters are energised by ATP hydrolysis, and the secondary active transporters utilise electrochemical gradients of protons or ions as their energy source ^7^. Further, the secondary active transporters are subdivided into several families, namely, the major facilitator superfamily (MFS), small multidrug resistance (SMR) family, resistance–nodulation–cell division (RND) family and the multidrug and toxic compound extrusion (MATE) family ^8^.

The MFS family of pumps are widely distributed amongst the mycobacterial species and are associated with antimicrobial resistance ^9^. These pumps utilise electrochemical gradients Proton Motive Force (PMF), and transport single polypeptides, toxic metabolites, heavy metals, and solvents across the cell membrane ^10^. LfrA, a MFS transporter, was the first reported efflux pump in *M. smegmatis* that confers low-level resistance to fluoroquinolones, ethidium bromide, acridine orange and some quaternary ammonium compounds ^11^. Tet38, a single peptide chromosome-encoded pump, is one of the reasons for the resistance of *S. aureus* towards tetracycline and fosfomycin ^12^. PMF-dependent Rv1877 of *Mtb* and MSMEG_2991 of *M. smegmatis,* are reported to exert low-level of resistance to multiple fluoroquinolones and unrelated classes of antibiotics ^13,14^. Pumps such as PMT belonging to the MFS class also play major roles in enhancing the biofilm synthesis of *Acinetobacter baumanii* ^15^. Several other pumps, such as EfpA, Tap, and P55, confer resistance to multiple unrelated, clinically relevant antimicrobial drugs ^16–18^.

Rv1250 is a probable MFS-type membrane-bound transporter protein belonging to *M. tuberculosis* (*Mtb*). It is reported that prolonged anti-tubercular drug therapy escalates the level of expression of Rv1250 ^19^. Generally, mutations in *katG* and *inhA* are attributed as the major reasons behind isoniazid resistance in *Mtb*. However, a high-level isoniazid-resistant isolate, with no mutations in *katG* and *inhA*, is reported to over-express Rv1250 ^20^. Another study reveals that among the mono-resistant isolates, Rv1250 and Rv876 are co-overexpressed in response to rifampicin, and in the presence of verapamil, there is a significant reduction of expression of Rv1250 and Rv876 ^21^.

Since Rv1250 of *Mtb* is found to be over-expressed upon anti-tubercular drug therapy, to find the rationale of such overexpression, here we intend to demonstrate the plausible role of Rv1250 in contributing to antimicrobial resistance, which is in form of extruding multiple structurally unrelated classes of drugs, boosting host biofilm forming ability and facilitating the cell survival.

## Results

### Phylogenetic analysis of Rv1250 homologs reveals an evolutionary conservation across pathogenic and environmental *Mycobacteria*

The phylogenetic tree created using the homologous protein sequence of Rv1250 from 19 *Mycobacterium* species employing the Maximum Likelihood method with 1000 bootstrap replicates and E-value < 1e−5, shows distinct clades, indicating potential differences in ecological niches or pathogenicity profiles. All three slow-growing pathogenic species, *M. tuberculosis, M. canettii,* and *M. decipiens*, cluster together, suggesting a conserved protein function of Rv1250 within the *M. tuberculosis* complex (MTBC). The same was seen in species like *M. avium* complex (MAC), known for causing pulmonary and disseminated diseases, particularly in immune-compromised individuals, and in *M. branderi*, *M. kyorinense*, and *M. celatum,* which belong to slow-growing non-tubercular mycobacteria (NTM). Similarly, it was also conserved in *M. riyadhense*, *M. bourgelatii*, *M. gordonae*, and *M. asiaticum,* described as environmental or opportunistic pathogens (**Figure 1**).

**Figure 1:**
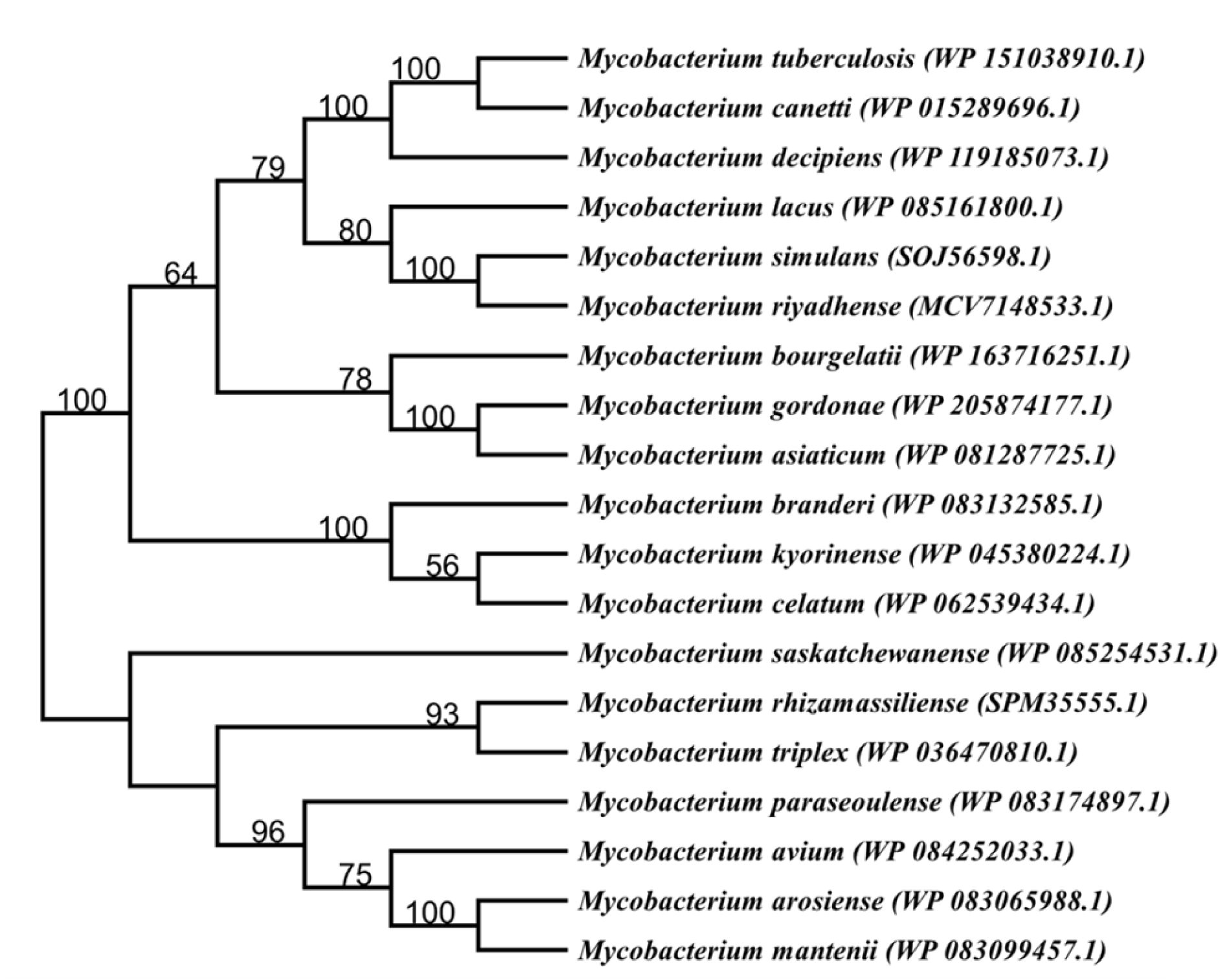
Phylogenetic tree analysis: Maximum Likelihood phylogenetic tree based on the Rv1250 protein sequence and its homologs (E-value < 1e−5) from representative Mycobacterium species. The tree was reconstructed with 1000 bootstrap replicates. Bootstrap values greater than 50% are indicated at the corresponding nodes.

### Expression of Rv1250 decreased the susceptibility of the host cells towards structurally unrelated classes of antibiotics

To determine the role of Rv1250 in influencing the antimicrobial activity of *E. coli* and *M. smegmatis*, the gene was expressed in the host cells (**Figure 2 and S1**), and the variation in susceptibility of the host towards different classes of antibiotics was studied. The expression of *rv1250* reduced the susceptibilities of the host cells towards the fluoroquinolone antibiotics tested, namely, norfloxacin (4 fold), ofloxacin (4 fold), and levofloxacin (2 fold) in comparison to the control cells harbouring the empty vector. Likewise, enhanced resistance towards beta-lactam antibiotics and aminoglycosides, ampicillin (4 fold) and oxacillin (4 fold), amikacin (2 fold), apramycin (2 fold) and gentamicin (2 fold) was also noted for *E. coli* and *M. smegmatis* cells (**Table S1 and Table 1**). Furthermore, *M. smegmatis* cells expressing *rv1250* showed resistance towards anti-tubercular drugs, *viz*., rifampicin (32 fold), ethambutol (4 fold), and isoniazid (16 fold) (**Table 1**). Additionally, antibiotic susceptibilities of the host cells harbouring Rv1250 were tested in the presence of a sub-inhibitory concentration of the PMF blocker, carbonyl cyanide m-chloro-phenylhydrazone (CCCP). It was observed that the presence of CCCP reduced the difference in the level of resistance between the control cells and the experimental cells towards all the tested antibiotics to 1 – 2 fold (**Table 1 and S1**). The reduction in the level of susceptibility of the host cells expressing Rv1250 in presence of CCCP indicated the involvement of the protein in imparting multidrug resistance, and CCCP might act as an inhibitor for Rv1250.

**Figure 2:**
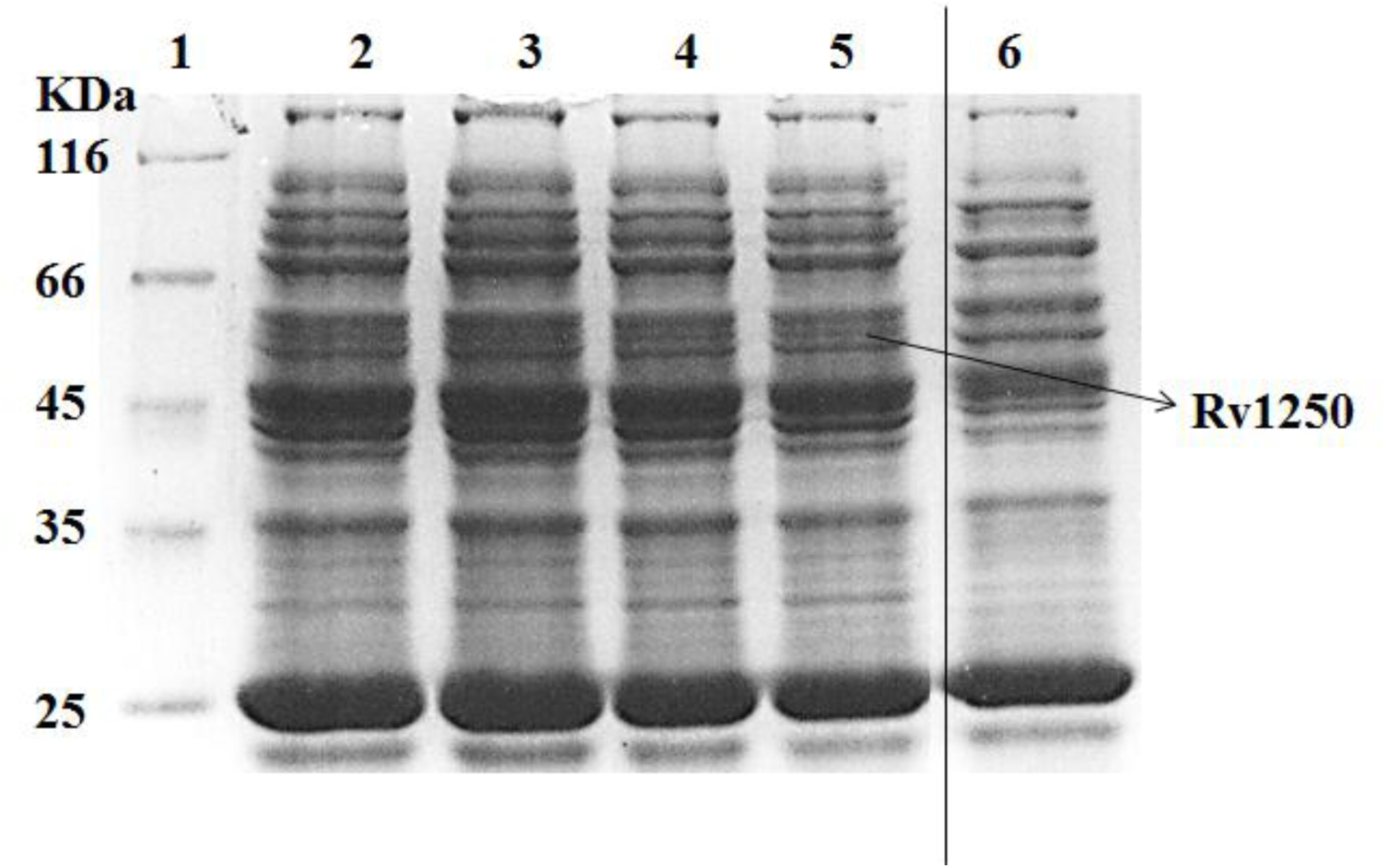
SDS-PAGE image of solubilized membrane fraction of *M. smegmatis* MC^2^155 cells harbouring pM1250 upon inducing with 20 ng ml^-1^ tetracycline: Lane1: unstained protein molecular weight marker (Thermo Scientific, Waltham, MA, US), Lane2: induced pM1250-Q409A in *M. smegmatis* cells, Lane3: induced pM1250-Q409E Lane4: induced pM1250-D34A, Lane5: *M. smegmatis* cells expressing pM1250, Lane 6: uninduced Rv1250 in *M. smegmatis* cells.

**Table 1:** Minimum Inhibitory Concentrations of various unrelated classes of antibiotics against *M. smegmatis* cells harbouring pMIND vector (control of *M. smegmatis* cells) and Rv1250 (pM1250) and its mutants in the presence and absence of CCCP.

| Antibiotics | Minimum Inhibitory Concentration (MIC) mg l<sup>-1</sup> |  |  |  |  |  |  |  |  |  |
| --- | --- | --- | --- | --- | --- | --- | --- | --- | --- | --- |
|  |  |  |  |  |  | CCCP |  |  |  |  |
|  | *M. smeg* | Rv1250 | Q409A | Q409E | D34A | *M. smeg* | Rv1250 | Q409A | Q409E | D34A |
| Norfloxacin | 2 | 8 | 16 | 2 | 2 | 1 | 2 | 2 | 1 | 1 |
| Ofloxacin | 0.5 | 2 | 4 | 0.5 | 0.5 | 0.25 | 0.25 | 0.25 | 0.25 | 0.25 |
| Levofloxacin | 2 | 4 | 8 | 2 | 2 | 1 | 2 | 2 | 1 | 1 |
| Ampicillin | 32 | 128 | 256 | 32 | 32 | 8 | 16 | 16 | 8 | 8 |
| Oxacillin | 16 | 64 | 128 | 16 | 16 | 4 | 4 | 4 | 4 | 4 |
| Amikacin | 4 | 8 | 16 | 4 | 4 | 2 | 2 | 2 | 2 | 2 |
| Apramycin | 4 | 8 | 16 | 4 | 4 | 4 | 4 | 4 | 4 | 4 |
| Gentamicin | 8 | 32 | 64 | 8 | 8 | 2 | 2 | 2 | 2 | 2 |
| Isoniazid | 32 | 512 | 1024 | 32 | 32 | 8 | 16 | 16 | 8 | 8 |
| Rifampicin | 1 | 32 | 64 | 1 | 1 | 0.5 | 8 | 16 | 0.5 | 0.5 |
| Ethambutol | 0.18 | 0.72 | 1.44 | 0.18 | 0.18 | 0.09 | 0.09 | 0.09 | 0.09 | 0.09 |

### *rv1250* reduced the intracellular antibiotic/dye accumulation levels in the host cells

It was observed that Rv1250-expressing cells showed reduced levels of susceptibility towards different classes of antibiotics, namely fluoroquinolones, beta-lactams, aminoglycosides and anti-tubercular drugs. There was no change in the growth pattern of control cells and *rv1250* expressing *M. smegmatis* cells (**Figure S2**). To ascertain the role of Rv1250 in antimicrobial resistance, intracellular antibiotic/dye accumulation and efflux assays were performed. It was observed that *E. coli* cells expressing Rv1250 accumulated 55% less norfloxacin (fluoroquinolone antibiotic) within 15 min as compared to control cells carrying empty pBAD-18Cam vector (**Figure S3**). Similarly, *M. smegmatis* cells harbouring *rv1250*, cloned in the pMIND vector and induced with 20 ng ml^-1^ of tetracycline, showed reduced level of accumulation of norfloxacin by 50% within 18 min (**Figure 3**). Likewise, the accumulation levels of Bocillin FL, a fluorescent penicillin (beta-lactam antibiotic), were also reduced by 40% within 15 min (**Figure 3**) ^22^. When, CCCP was introduced into the experimental setup ^23^, led to an increase in accumulation levels of the tested antibiotics in both the control as well as the Rv1250 expressing experimental cells. The observed change was estimated by the rapid increase in the fluorescence signal (**Figures S3 and 3**) leading to the levels of accumulation between the control cells and the experimental cells to near equal levels. These results show that *rv1250* increases the resistance of the host cells towards various antibiotics by reducing the intracellular accumulation levels, which could be nullified in the presence of an inhibitory effect of CCCP.

**Figure 3:**
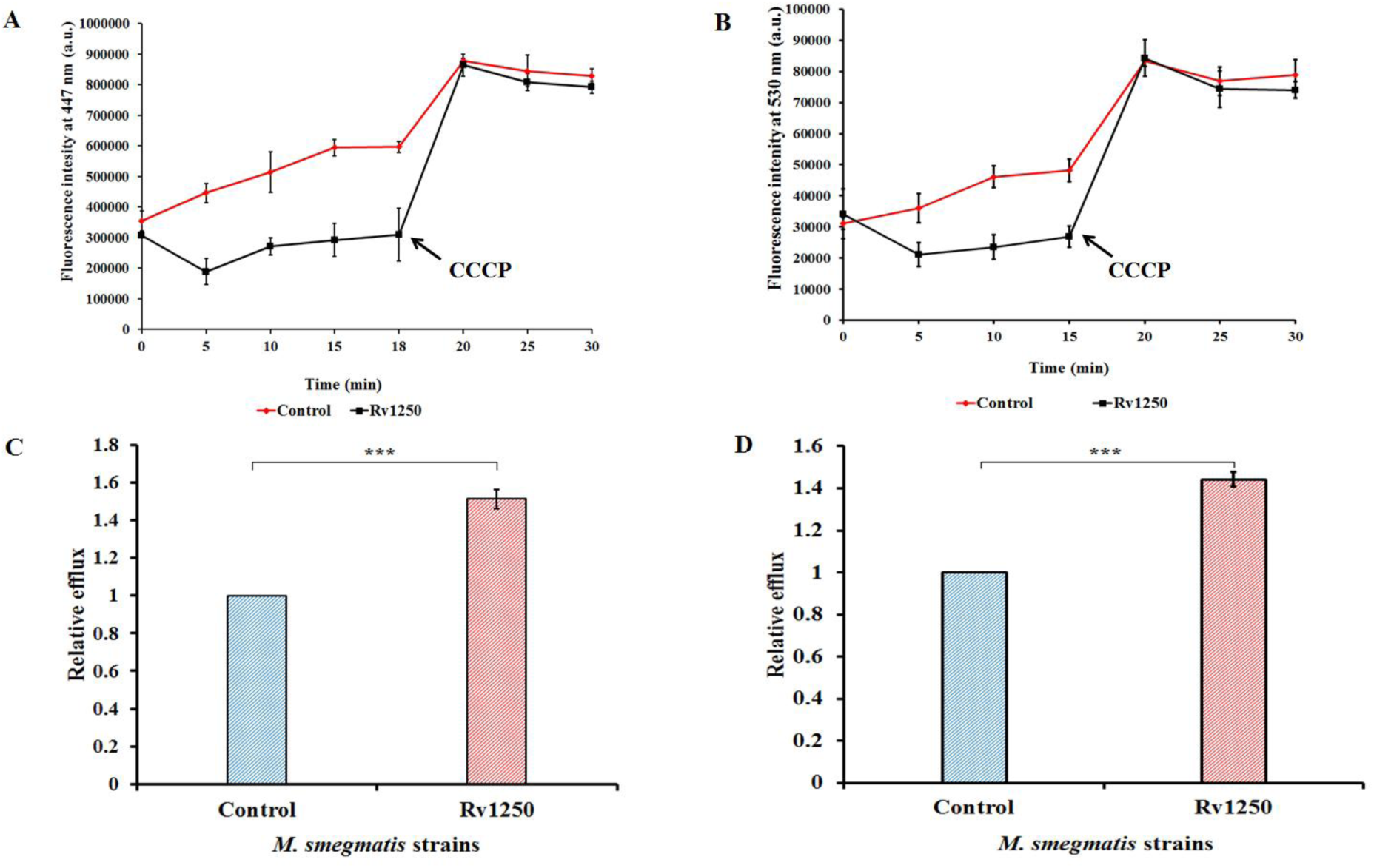
Intracellular Norfloxacin and Bocillin FL accumulation assay in *M. smegmatis*: Accumulation of norfloxacin (A) and Bocillin FL (b) by the cells harbouring Rv1250 in comparison to control cells (harbouring empty vector) with respect to time. The antibiotics were added at the 0^th^ min. Sample aliquots were drawn at every 5 min interval for a period of 30 min, and CCCP was added at 15^th^ - 18^th^ min. Relative efflux was calculated after exposing the *M. smegmatis* to norfloxacin (C) and Bocillin FL (D) cells for 5 – 10 min.

On the other hand, Rv1250 expressing *M. smegmatis* cells also enhanced the efflux of EtBr. Real-time EtBr efflux assay was performed in the presence of CCCP. The initial level of fluorescence of the preloaded dye was similar for both the control and the experimental cells, which were recorded for ∼50 sec. However, upon energizing the cells with glucose, a rapid decrease (∼ 60%) in the fluorescence intensity was observed in the host cells expressing Rv1250 than that of control cells (**Figure 4**). The cells expressing *rv1250* extruded ∼ 60 % more EtBr in comparison to the cells harboring empty vector after energization of the cells with glucose. Therefore, the observations indicate the role of the active efflux pump Rv1250 in imparting resistance towards toxic dyes like EtBr.

**Figure 4:**
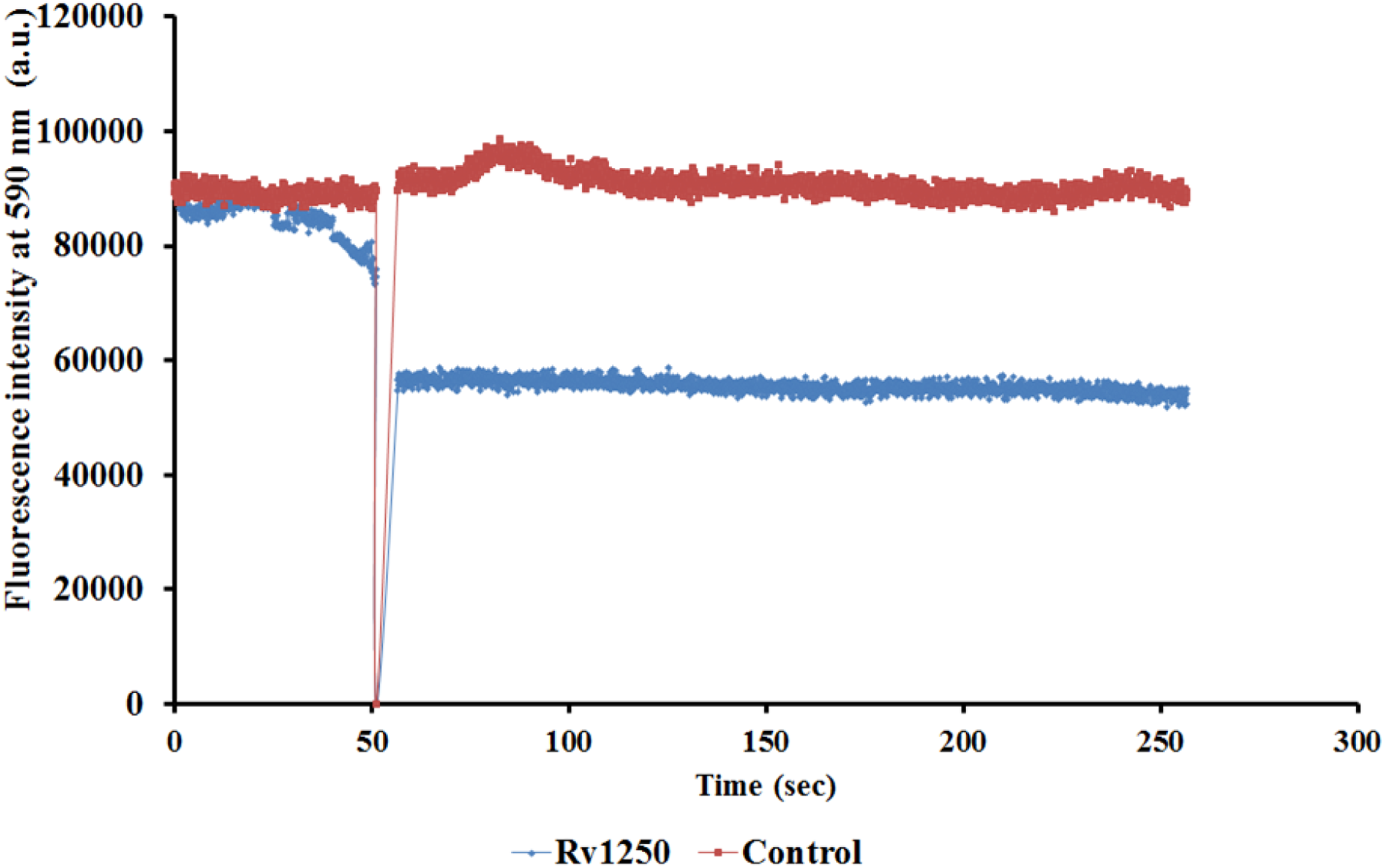
Real-time EtBr efflux assay: EtBr efflux from *M. smegmatis* cells harbouring Rv1250 and empty vector control cells devoid of Rv1250, after energisation with glucose at ∼50 sec for 200 sec.

### The substitutions of Q409E, Q409A and D34A affect the efflux activity of Rv1250

It was previously hypothesised by Heng *et al*. that D34, is a residue of MFS-type transporter that is involved in proton and substrate transport ^24^. To study the effect of the residue D34 of Rv1250 on the antimicrobial efflux activity of the host cells expressing *rv1250*, site-directed mutagenesis was performed. Single amino acid substitution (alanine) of residue D34 led to a reduction of the efflux activity of Rv1250. The mutation created a complete loss of resistance of the host cells towards fluoroquinolones, aminoglycosides, beta-lactams and anti-tubercular drugs (**Table 1 and S1**). Further, the cells expressing Rv1250_D34A_ increased the accumulation of norfloxacin and Bocillin FL in the host cells (**Figure 5**).

**Figure 5:**
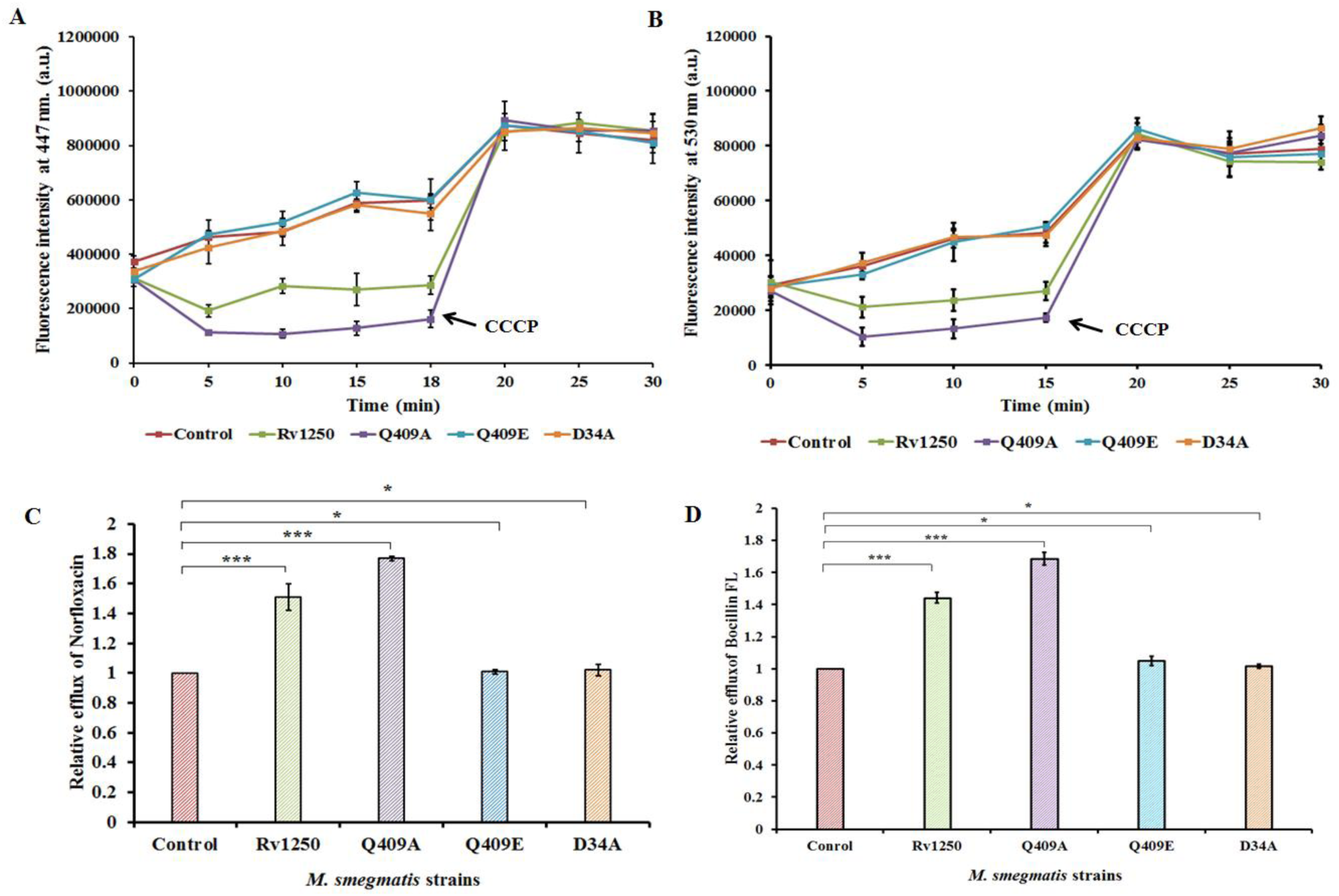
Intracellular Norfloxacin and Bocillin FL accumulation assay in *M. smegmatis* expressing *rv1250* and its mutants: Accumulation of norfloxacin (A) and Bocillin FL (b) by the cells harbouring Rv1250 and its mutants in comparison to control cells (harbouring empty vector) with respect to time. The antibiotics were added at the 0^th^ min. Sample aliquots were drawn at every 5 min interval for a period of 30 min, and CCCP was added at 15^th^ - 18^th^ min. Relative efflux was calculated after exposing the *M. smegmatis* to norfloxacin (C) and Bocillin FL (D) cells for 5 – 10 min.

Molecular docking studies of Rv1250 with isoniazid suggested that Q409 may be critical in the transport mechanism. To prove this, we introduced site-directed mutations at Q409 — substituting glutamine with alanine (Q409A) to neutralise its effect, and with glutamic acid (Q409E) to alter its charge. It was observed that when the residue Q409 was substituted with glutamic acid a complete loss of the efflux activity of the host cells expressing the mutated Rv1250 was observed in comparison to the cells expressing wild-type Rv1250 (**Figure 5**). On the other hand, upon substitution of Q409 with alanine, the resistance of the host cells was increased by 2 – 4 fold than that of the cells expressing wild-type Rv1250. Taken together, it can be inferred that the residues Q409 and D34 are important for the antibiotic efflux activity and might play significant roles in maintaining the susceptibility pattern of host cells expressing Rv1250.

### Rv1250 enhances the biofilm formation of the host cells

Efflux pumps have been shown to play a critical role in biofilm formation ^25^. They are often linked with the release of extracellular polymeric substances (EPS) and quorum-sensing molecules, which contribute to the survival of the microorganism in adverse conditions ^26^. Therefore, to understand the effect of *rv1250* on the biofilm formation of the host cells, a 24-well plate static biofilm formation assay was performed. A 2 – 4 fold increase in biofilm formation by the host cells expressing Rv1250 was observed than that of control cells (**Figure 6 and S4**). In the presence of 3 µg ml^-1^ of CCCP, an uncoupler of oxidative phosphorylation which disrupts the ionic gradient of bacterial membranes and blocks efflux pumps, the difference in the biofilm-forming capacity of the experimental cells and the control cells was reduced to less than 2-fold (**Figure 6**) ^23^. Further, to determine difference in the amount of carbohydrate and protein content in the biofilm matrices of the experimental cells and control cells the chloroform: methanol extraction method was performed. It indicated an increase in protein and carbohydrate concentration in the biofilm matrices of the *rv1250* expressing cells by 28 ± 0.32 % and 46.01 ± 0.76 %, respectively, in comparison to the host *M. smegmatis* cells containing of empty pMIND. Likewise, the biofilm-forming abilities of the host cells expressing the mutated Rv1250_D34A_ and Rv1250_Q409E_ were also reduced as compared to wild-type Rv1250-expressing cells (**Figure 7**). On the other hand, the *M. smegmatis* cells expressing the Q409A mutant showed an increase in biofilm formation of the host cells by more than 4-fold. Therefore, it was observed that Rv1250 contributed in enhancing the biofilm formation of the host cells. The mutants expressing Rv1250_D34A_ and Rv1250_Q409E_ reduced and Rv1250_Q409A_ increased the biofilm forming ability of the host cells just as they affected the efflux activity of Rv1250.

**Figure 6:**
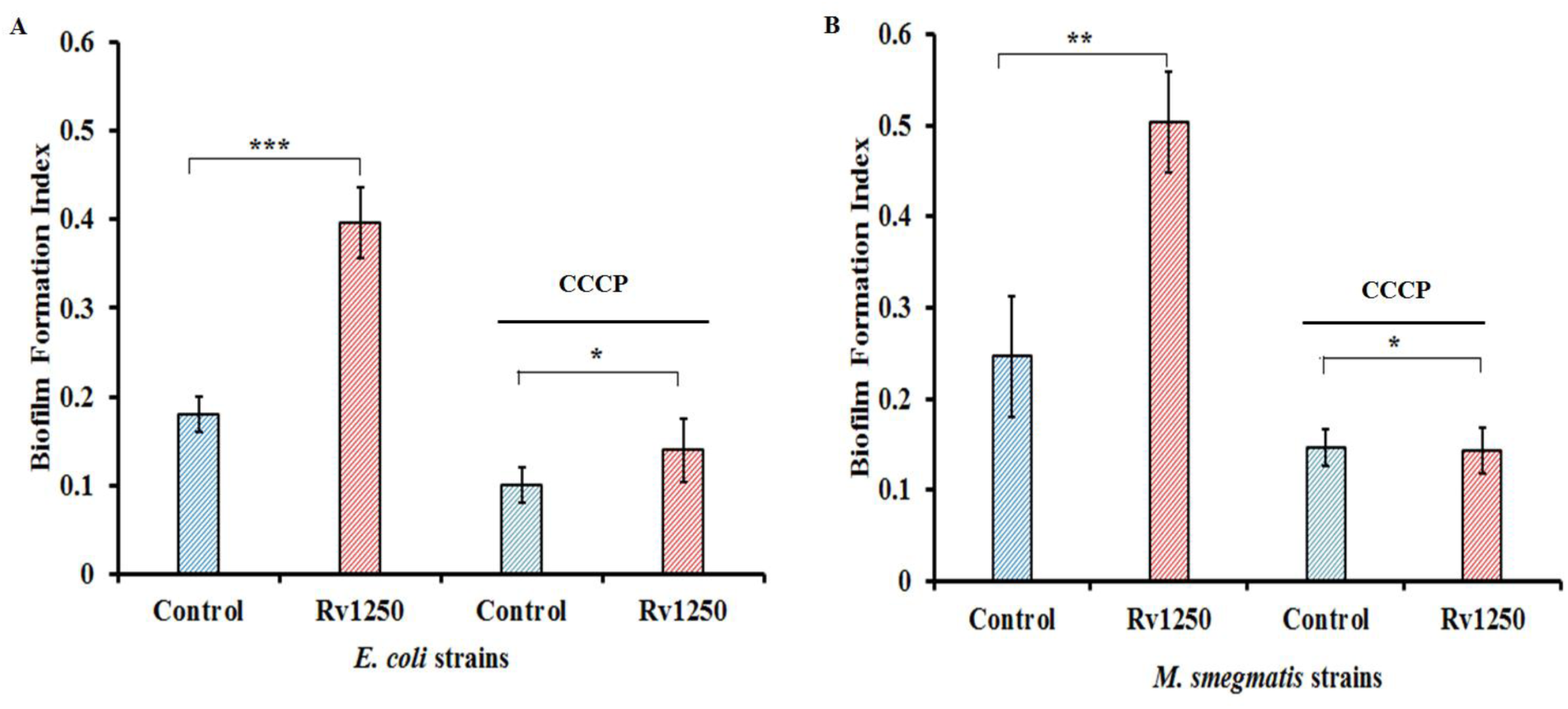
Semi-Quantitative biofilm formation assay: Biofilm formation by *E. coli* CS1O9 harbouring Rv1250 (A), *M. smegmatis* cells expressing Rv1250 and empty vector control cells (B). Biofilm quantification in the presence of CCCP by crystal violet staining assay in *rv1250* expressing *E. coli* (A) cells and *M. smegmatis* (B) cells, respectively.

**Figure 7:**
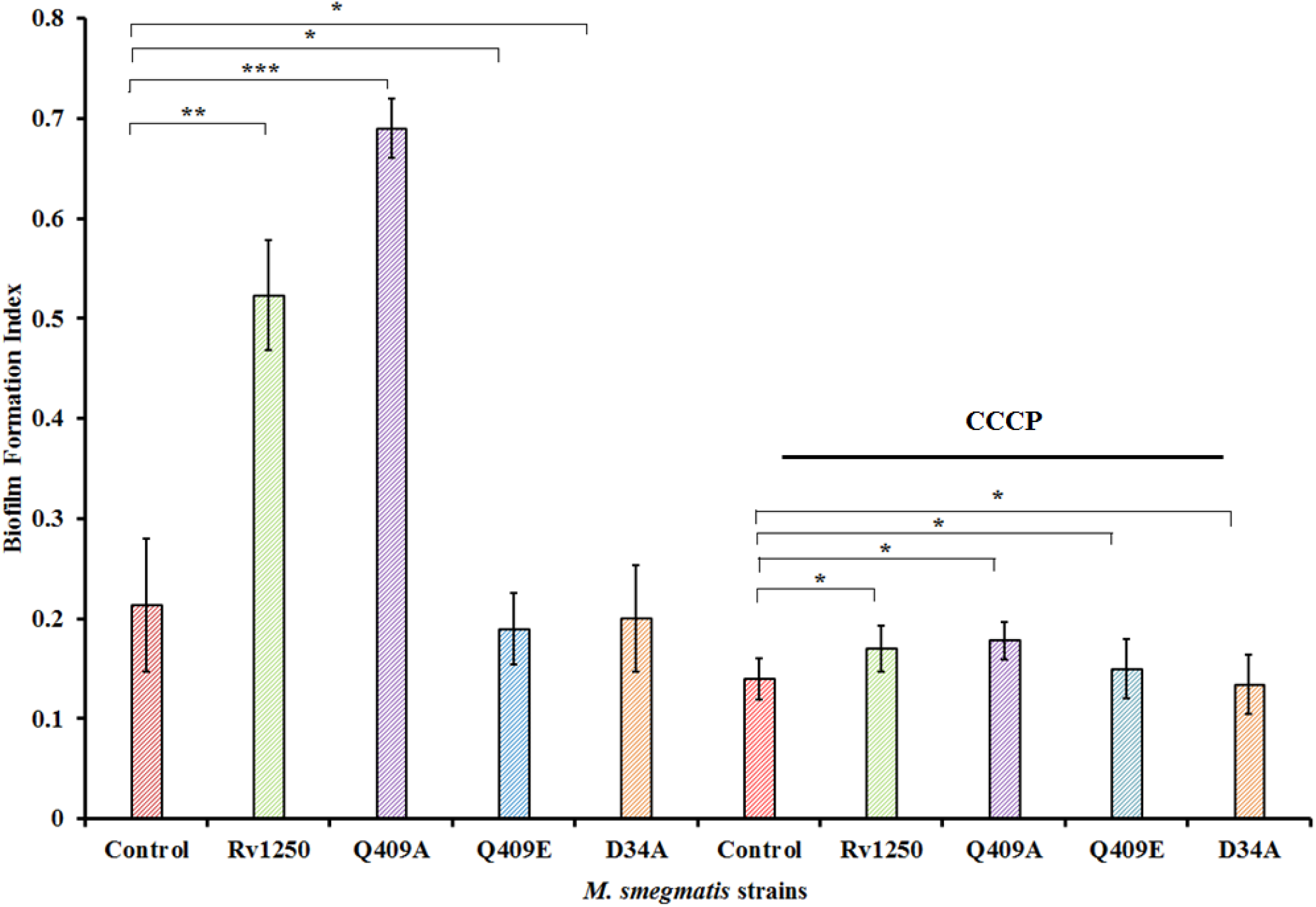
Semi-Quantitative biofilm formation assay of *rv1250* and its mutants expressing *M. smegmatis* cells: Biofilm formation by *M. smegmatis* cells expressing Rv1250, its mutants and empty vector control cells in the presence and absence of CCCP.

### Rv1250 enhances mycobacterial infection in RAW 264.7

Mycobacteria are known to survive and proliferate in host macrophages, combating antimicrobial resistance ^27^. Mycobacterial efflux pumps and their regulators were shown to play a role in macrophage infection and survival ^28–31^. Therefore, to demonstrate the role of Rv1250 in macrophage infection and survival, survival studies of *M. smegmatis* cells in RAW 264.7 were conducted. It was observed that in the presence of sub-inhibitory concentrations of rifampicin (0.1 mg l^-1^) an anti-tubercular drug, the *M. smegmatis* cells expressing *rv1250* were survived more within the RAW 264.7 cells in comparison to the cells carrying the empty pMIND (**Figure 8B)**. Further, when CCCP was introduced into this experimental set-up, the relative difference in survival percentage of both the *M. smegmatis* cells harboring empty pMIND vector *and M. smegmatis* cells expressing *rv1250* reduced. The relative survival percentage of all the cells were similar after addition of CCCP (**Figure 8C**), indicating an inhibitory effect of CCCP on Rv1250. However, in the absence of rifampicin, there was no change in the relative survival percentage of the Rv1250 expressing *M. smegmatis* cells and the cells carrying empty pMIND (**Figure 8A**).

**Figure 8:**
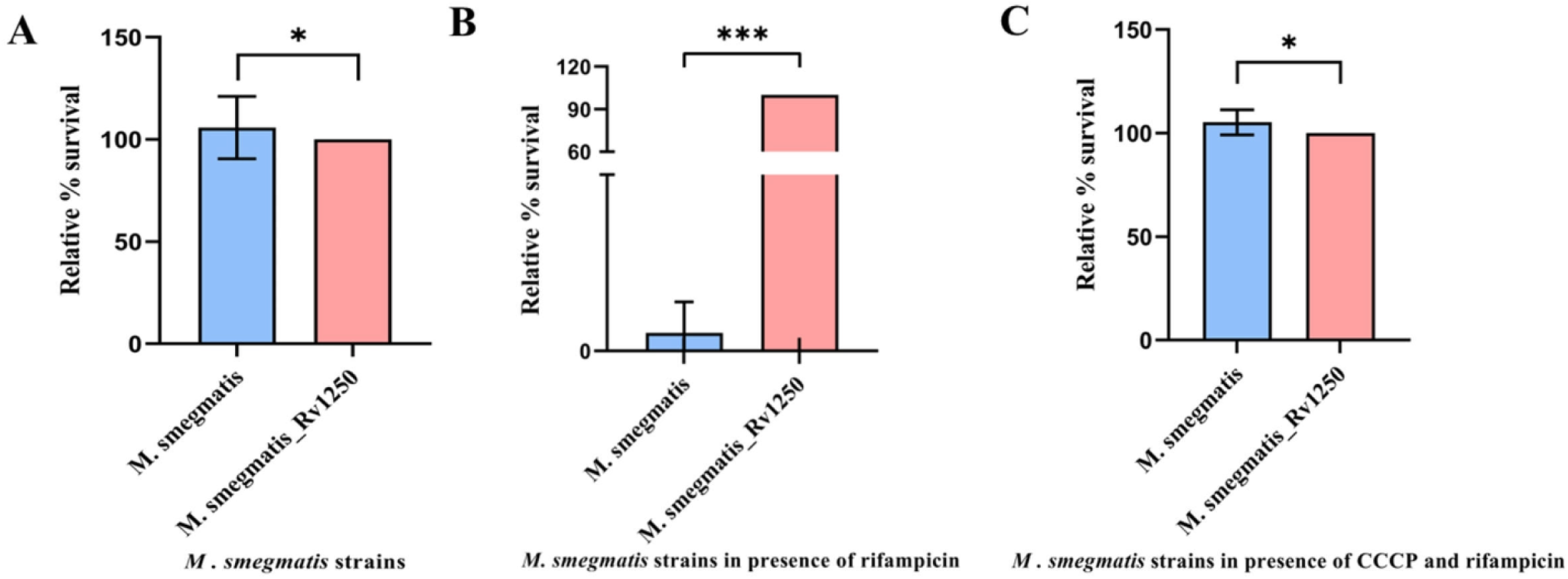
Rv1250 confers tolerance within Macrophage: Relative percentage survival of bacterial cells inside RAW 264.7 (A), Relative percentage survival of bacterial cells inside RAW 264.7 in presence of sub inhibitory concentration of rifampicin (B), Relative percentage survival of bacterial cells inside RAW 264.7 in presence of sub inhibitory concentration of rifampicin and CCCP (C).

Therefore, it was speculated that Rv1250 plays a role in assisting the survival of *M. smegmatis* cells in macrophages in the presence of rifampicin.

### MD simulation revealed differential ligand binding in the central cavity of Rv1250 and its mutants

Rv1250 is a putative Major Facilitator Superfamily (MFS) transporter protein in *Mycobacterium tuberculosis*. Our *in vivo* data indicate its involvement in antimicrobial efflux. To elucidate the molecular mechanism underlying this function, we performed molecular docking of Rv1250 with isoniazid, followed by a 500 ns all-atom molecular dynamics (MD) simulation. Our analysis suggests that Q409 may be critical in the transport mechanism. To prove this, we introduced site-directed mutations at Q409 — substituting glutamine with alanine (Q409A) to neutralise its effect, and with glutamic acid (Q409E) to alter its charge. In addition, D34, a residue previously implicated in MFS transporter function, was mutated to alanine (D34A) ^24^. As noted, Q409E and D34A mutations abolished Rv1250 efflux activity, whereas Q409A significantly enhanced export.

Subsequently, 500 ns MD simulations were performed for each mutant-protein complex. All systems reached structural stability after 150 ns, with root mean square deviation (RMSD) values within 1 Å (0.1 nm). Radius of gyration (Rg) values remained stable throughout the simulations, supporting overall structural integrity. A minor RMSD deviation within 1 Å was observed for the D34A mutant between 200–400 ns. Notably, all mutants exhibited increased flexibility in residues 442–447, as indicated by elevated root mean square fluctuation (RMSF) values exceeding 1 Å relative to the wild-type complex (**Figure S5**). Despite these localised dynamics, no substantial global structural deviations were detected. However, significant differences emerged in ligand behaviour within the central cavity of the mutant protein-ligand complexes compared to the wild type. Principal component analysis (PCA), based on protein backbone and ligand atoms, and free energy landscape (FEL) analysis were conducted to characterise the ligand’s conformational sampling (**Figure 9C**).

**Figure 9:**
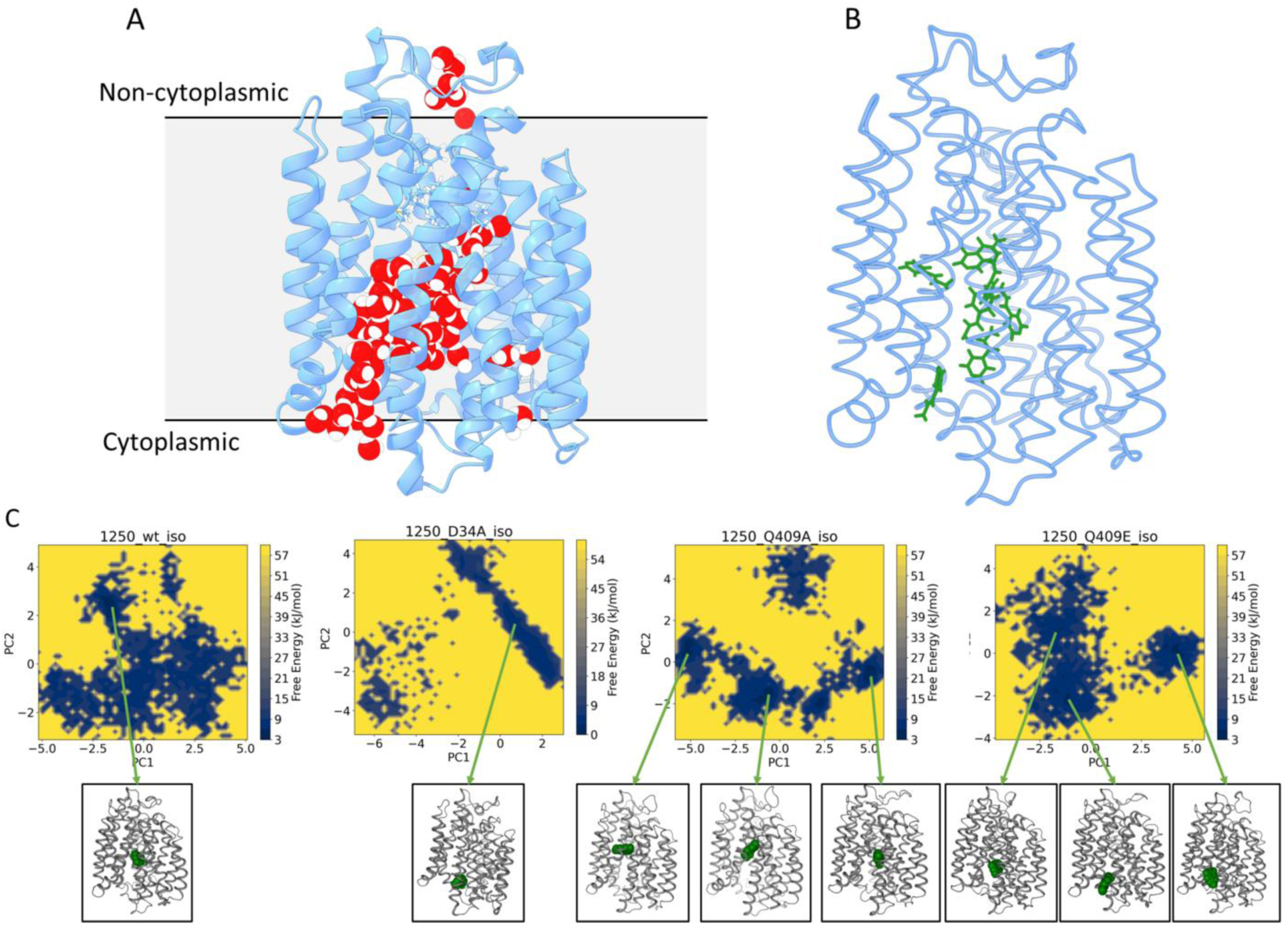
***In Silico* Analysis of Rv1250:** (A) Representative snapshot from the 500 ns molecular dynamics (MD) simulation of the native Rv1250-isoniazid complex, illustrating the water-filled central cavity. The grey background indicates the approximate position of the membrane. (B) Overlay of various isoniazid conformers visualized on the initial frame of the native protein-ligand complex. (C) Free energy landscapes and corresponding low-energy conformations of Rv1250-isoniazid complexes (wild-type, D43A, Q409A, and Q409E mutants) were obtained from the MD trajectories’ principal component analysis (PCA). The protein is shown in tube representation, and the isoniazid molecule is depicted as van der Waals spheres. For the wild-type protein, multiple low-energy conformations were sampled; however, only the conformation closest to the initial docked pose is presented.

In the wild-type complex, isoniazid explored multiple binding sites (**Figure 9B**) within the central cavity (**Figure 9A**), deviating from its initial docked position, with the FEL revealing several low-energy conformations indicative of dynamic ligand binding. In contrast, the D34A mutant facilitated rapid ligand migration toward the cytoplasmic end, where it remained stably bound, corresponding to a low-energy conformation (**Figure 9C**). Mutation of Q409 to alanine promoted ligand sampling closer to the extracytoplasmic side in energetically favourable poses. In contrast, substitution with glutamic acid restricted the ligand’s low-energy conformations toward the cytoplasmic side (**Figure 9C**).

As an efflux transporter, Rv1250 is proposed to export antibiotics from the cytoplasmic side to the extracellular environment. Therefore, it is plausible that substrates adopting a stable conformation closer to the extracellular side along the transport pathway may facilitate more efficient export. This mechanism may account for the differential transport behaviours observed in the Rv1250 mutants in our *in vivo* studies.

## Discussion

Efflux pumps are one of the most important and latest mechanisms identified to be involved in anti-microbial resistance ^32^. These pumps help regulate the internal environment of the microorganism by extruding toxins, metabolites, drugs, dyes, and quorum-sensing molecules ^33^. The conservation of Rv1250 across different mycobacterial species groups, including MTBC,

MAC, and NTM, emphasizes its vital role in the bacterium’s survival by transporting toxic compounds outside the cell. Few available reports suggest that Rv1250, a putative uncharacterized MFS-type transporter, might play a role in imparting antimicrobial resistance ^34–36^. However, very little experimental evidence is present now of the antimicrobial efflux activity of Rv1250. Therefore, in this present study, we demonstrate the role of Rv1250 in *M. tuberculosis* as a multidrug efflux pump in *E. coli* and *M. smegmatis*.

MFS-type secondary active transporters are one of the largest classes of efflux pumps known to regulate the cellular environment by extruding single polypeptides, toxic metabolites, heavy metals, and solvents ^10,37^. Expression of Rv1250 in *E. coli* and *M. smegmatis* cells decreased the susceptibility of the host cells towards an array of structurally unrelated classes of antibiotics, namely fluoroquinolones, beta-lactams, aminoglycosides and anti-tubercular drugs. These antibiotics are known to play key roles as first or second line of drugs needed for tuberculosis treatments. Due to the present findings we suggested that Rv1250 might contribute to providing low to moderate level intrinsic tolerance towards a wide range of substrates. Therefore, to further establish the underlying reason behind the increase in tolerance of the Rv1250 expressing cells towards a multitude of antimicrobials, we performed intracellular antibiotic accumulation assays and real-time efflux assays.

Expression of Rv1250 under arabinose and tetracycline-inducible promoters led to a considerable decrease in accumulation levels of fluoroquinolone and beta-lactams inside the cells. To further verify its energy source, a PMF blocker, CCCP, known to inhibit pumps that utilise PMF as their energy source, was incorporated into the experimental setup ^38^. CCCP increased the accumulation levels of antibiotics in both the control cells and the experimental cells. The difference in the accumulation levels between the control and experimental cells became almost negligible, indicating the inhibitory effect of CCCP on *rv1250*. Henceforth, it is observed from the present study that Rv1250 is a secondary active efflux pump energized by PMF and extrudes a variety of toxic compounds out of the cells rendering it resistant to antimicrobials.

EtBr, a toxic fluorescent dye, is often used as a substrate to determine the efflux pump activity of cells ^39^. EtBr emits weak fluorescence in an aqueous environment and strongly fluoresces when present in the periplasm of Gram-negative bacteria and the cytoplasm of Gram-positive bacteria ^40^. Therefore, EtBr was utilised to study the efflux activity of Rv1250-expressing cells. It is observed that the levels of EtBr in the preloaded cells were similar for the control and the experimental cells. However, energisation with glucose led to rapid efflux of EtBr from the Rv1250 expressing cells in comparison to the control cells consisting of an empty vector. This observation suggests that Rv1250 actively extrudes toxic dyes like EtBr out of the cell.

The residues D34 and Q409 that line the central pore of the transporter are subjected to site-directed mutagenesis. Upon mutating the D34 residue to alanine, there was complete abolishment of the efflux activity of Rv1250. It was previously hypothesised by Heng *et al*. that D34, is a residue of MFS-type transporter that is involved in proton and substrate transport ^24^. Further, according to the present findings, it has been observed that upon mutating aspartic acid to alanine at the 34^th^ position, there is an interruption in the migration of the antibiotics outside the cell. It is speculated that a stable complex between the pump and the substrate is formed that leads to the abolishment of transporter activity of Rv1250. Similarly, the 409^th^ amino acid residue is mutated to alanine. Upon mutating glutamine to alanine, the transport activity of Rv1250 is enhanced. On the other hand, mutating glutamine to glutamic acid led to a complete loss of efflux activity of Rv1250. It is speculated that upon mutating the 409^th^ residue of Rv1250 to alanine, it promoted the substrate (antibiotic) movement more towards the extracytoplasmic side by facilitating energetically favourable substrate protein binding conformations. On the other hand, upon mutating glutamine to glutamic acid, there is a complete loss of efflux activity due to constrained low-energy interactions towards the cytoplasmic region. Therefore, it could be contemplated that D34 and Q409 are important residues that facilitate the efflux activity of Rv1250.

Efflux pumps facilitate biofilm formation by transporting nutrients, assisting cells in extruding toxins, metabolites, and protecting cells from pH, salt, and antibiotic stress ^25,26^. Multiple efflux pumps are known to enhance the biofilm formation of the host cells ^13,14,41–43^. It was also observed that *rv1250,* like multiple known efflux pumps, enhanced the biofilm-forming capacity of the host cells by 2 – 4 fold. There was an increase in carbohydrate and protein concentration in the biofilm matrices of the host *M. smegmatis* cells expressing *rv1250* in comparison to empty pMIND vector consisting of *M. smegmatis* cells, which might result increase in biofilm formation ^44^. Biofilm maturation increases the carbohydrate and protein concentration within the matrices which leads to an increased biofilm density and a robust core ^45^. Further to confirm that indeed *rv1250* is involved in influencing the biofilm formation of cells expressing it, CCCP was used in the 24-well plate static biofilm quantification assay ^23^. The addition of CCCP reduced the difference in BFI between the control and the experimental cells to less than 2-fold. Therefore, this indicates that Rv1250 is involved in influencing the biofilm-forming capability of the host cells expressing it along with extruding multiple structurally unrelated classes of toxins.

Apart from enhancing the drug efflux activity of the cells, the pumps also contribute to biofilm formation, morphology, virulence, cell wall assembly, and growth ^46–49^. In this study, it is observed that in the presence of sub-inhibitory concentrations of rifampicin, there is a significant difference in the survival of the cells within macrophages. The cells expressing Rv1250 substantially survived more than the *M. smegmatis* cells devoid of Rv1250. In the presence of CCCP, the survival percentage of all the tested *M. smegmatis* cells was similar. Again, in the absence of any external antimicrobial stress, the survival percentage of all the bacterial cells did not show a significant difference. Hence, it could be speculated that Rv1250 provides tolerance to the cells expressing it towards anti-tubercular drugs like rifampicin. Upon addition of a sub-inhibitory concentration of CCCP, the difference in survival percentages of the tested bacterial cells was negligible, indicating the inhibitory effect of CCCP on Rv1250. It is speculated that Rv1250 is required for survival of *Mycobacterium smegmatis* within the RAW cells under antibiotic stress (rifampicin). Rv1250 extrudes anti-tubercular drug like rifampicin and reduces its concentration to sub-inhibitory levels thereby decreasing the effect of it on the microorganism.

MFS transporters facilitate substrate translocation across membranes via a rocker-switch mechanism, cycling through distinct conformations: inward-open, inward-occluded, occluded, outward-occluded, and outward-open. The PMF typically drives these transitions ^50^. Structural analysis of our model suggests that the transporter adopts an inward-occluded conformation. Notably, within the 500 ns MD simulation, no major conformational transitions were observed. This is likely due to the high energy barriers between states, which are difficult to overcome within the simulation timescale, particularly in the absence of an explicitly modelled PMF. To further explore substrate transport, umbrella sampling and extended adaptive biasing force (eABF) simulations using the z-axis as the collective variable was performed. However, these enhanced sampling approaches also failed to induce any significant conformational changes in the protein (data not shown), further supporting the notion of a substantial energy barrier for conformational transitions. The simulation study provides important insights into ligand behaviour within the central cavity, a characteristic feature of MFS transporters. The observed substrate behaviour may help explain the protein’s functional activity *in vivo*. While longer simulations or other enhanced sampling methods may be required to capture the full conformational cycle, our findings highlight key aspects of substrate binding and stability within the inward-occluded state.

## Conclusion

Rv1250 of *M. tuberculosis* plays an important role in enhancing the multidrug efflux activity of the studied experimental cells, *E. coli* and *M. smegmatis*. It helps in extruding multiple structurally unrelated classes of drugs and dyes. It also helps in enhancing biofilm formation. D34 and Q409 are important residues that facilitate the functioning of Rv1250. Rv1250 also facilitates in survival of mycobacteria in macrophages in the presence of rifampicin.

## Materials and Methods

### Bacterial strains, Plasmids, culture media and chemicals

The bacterial strains used in this investigation were *E. coli* XL1-Blue (*recA1 endA1 gyrA96 thi-1 hsdR17 supE44 relA1 lac*) (Stratagene/ Agilent, Santa Clara, CA, USA), *E. coli* CS1O9 (W1485 *glnV rpoS rph*) and *Mycobacterium smegmatis* MC^2^ 155 (ATCC). The *E. coli* strains were cultured at 37°C in Luria–Bertani broth (LB) or LB agar (Hi-Media, Mumbai, MH, India) with appropriate antibiotics (12 ng ml^-1^ tetracycline and 20 μg ml^-1^ chloramphenicol were used to grow *E. coli* XL1-Blue cells harbouring pBAD18-Cam vector) ^51^. *Mycobacterium smegmatis* MC^2^ 155 strains were cultured in Middle Brook 7H9 broth and 7H11 agar medium, (Sigma-Aldrich, St. Louis, MO, USA) supplemented with oleic acid-ADC enrichment (Hi-Media, Mumbai, MH, India), 0.35% (w/v) glycerol, 0.05% (w/v) Tween 80 and appropriate antibiotics (50 μg ml^-1^ of kanamycin and 100 μg ml^-1^ of hygromycin for the strains carrying pMIND and pSMT-100 vectors) ^52,53^. *E. coli* and *M. smegmatis* strains were tested for antibiotic susceptibility using cation-adjusted Mueller–Hinton Broth (MH) and 7H9 (HiMedia, Mumbai, MH, India), respectively. Unless otherwise specified, all the other reagents, including antibiotics, were purchased from Sigma-Aldrich (St. Louis, MO, USA).

### Construction of recombinant plasmids for *in vivo* studies

Using the primers for pMIND FP: 5’ CTC TCT CAT ATG AGG AGG CTC TCT CTA TGA CTA CGG CGA TAC 3’ and RP: 5’ CTC TCT ACT AGT TTA TAG TGC GCT CGG AGC TGT TGA CTC AGT ATC 3’ and, for pBAD-18Cam FP: 5’ CTC TCT GCT AGC AGG AGG CTC TCT CTA TGA CTA CGG CGA TAC 3’ and RP: 5’ CTC TCT AAG CTT TTA TAG TGC GCT CGG AGC TGT TGA CTC AGT ATC 3’, *rv1250* was amplified from the genomic DNA of *Mycobacterium tuberculosis* H37Rv. The amplicons were cloned at N*de*I-S*pe*I sites in the pMIND vector and N*he*I-H*ind*III sites in pBAD-18Cam vector, to construct pD1250 and pM1250, respectively ^51^. *E. coli* XL1-Blue and CS1O9 cells were transformed with pD1250 by heat shock, and pM1250 was electroporated into *Mycobacterium smegmatis* MC^2^155. The plasmid pD1250 was expressed in the CS1O9 by inducing the cells with 0.2% (w/v) arabinose. On the other hand, *rv1250* was expressed from pM1250 by inducing the *M. smegmatis* cells with 20 ng ml^-1^ tetracycline ^53^.

### Phylogenetic tree preparation

Homologous sequences were retrieved using BLASTP from the NCBI non-redundant (Nr) protein database, using the *Mycobacterium tuberculosis* Rv1250 protein sequence as the query. Only sequences with 100% query coverage and E-values less than 1e-5 were selected. Sequences with clearly defined species annotations were retained for downstream analysis. Multiple sequence alignment (MSA) was performed using Clustal Omega. A phylogenetic tree was constructed in MEGA12 using the Maximum Likelihood method based on the Jones-Taylor-Thornton (JTT) model of amino acid substitution ^54,55^. The tree with the highest log-likelihood score (−12,037.50) was selected for visualization. Bootstrap analysis was performed with 1,000 replicates.

### Expression of probable drug-transport integral membrane protein Rv1250 in *M. smegmatis*

Using 0.1% of the overnight cultured sample 10 ml of 7H9 broth was inoculated and incubated at 37°C with a shaking speed of 150 rpm till the culture reached an OD_600_ of ∼0.2. The *M. smegmatis* cultures were induced with 20 ng ml^-1^ of tetracycline to facilitate gene expression and protein synthesis, and incubated for 24 hr. The cells were spun down at 50,000 x g for 20 min at 4°C using a Sorvall RC6 PLUS centrifuge (Thermo Scientific, Waltham, MA, USA). The supernatant was discarded, and the resulting pellet was washed with 1 ml of 10mM Tris-Cl buffer (pH 7.5) and resuspended in the same buffer. The protease inhibitor, phenylmethylsulfonyl fluoride (PMSF), was added to the cell suspension at a final concentration of 1 mM. The cell suspension was sonicated in ice with five pulses of 60 sec each, followed by centrifugation at 50,000 x g for 10 mins. The supernatant fraction was collected and further centrifuged at 4°C for 1 hr. at 50,000 x g. The supernatant was discarded, and the pellet fraction was subsequently resuspended in 100 µl of Tris-Cl buffer (pH 7.5). The pellet fraction was further treated with 2% sarcosyl (sodium lauroyl sarcosinate) (w/v), mixed thoroughly, and incubated at 37°C on a thermomixer (Eppendorf, Hamburg, Germany) with shaking for 1 hr. to solubilise the proteins. The sample was centrifuged at 50,000 x g for 1 hr. at 4°C, and the supernatant was collected, estimated the protein concentration and finally analysed through sodium dodecyl sulphate polyacrylamide gel electrophoresis (SDS-PAGE) (12% acrylamide) ^53^.

### Assay for testing antimicrobial susceptibility of different classes of drugs and dyes

The minimum inhibitory concentration of multiple unrelated classes of drugs and dyes, such as fluoroquinolones, aminoglycosides, anti-tubercular drugs, beta-lactams, EtBr, acriflavine, and rhodamine B, was determined by a two-fold micro broth dilution method. The drugs and dyes were two-fold serially diluted in a 96-well plate, and the total volume was made up to 300 µl with MH broth containing 0.2% arabinose and 200 µl and with 7H9 media containing 20 ng ml^-1^ tetracycline to induce to induce rv1250 in *E. coli* and *M. smegmatis* cells, respectively ^41^. The plates were inoculated with bacterial strains (10^5^ per well) and incubated at 37°C. The optical density (OD_600nm_) of the culture medium was assessed after 12-16 hr. and 42-72 hr., respectively, for *E. coli* and *M. smegmatis* to assess the antimicrobial sensitivity as per CLSI (Clinical and Laboratory Standards Institute) guidelines by using a micro-plate reader (Biorad iMark™ Microplate Absorbance Reader, United States) ^56^. Further, CCCP at a sub-inhibitory concentration (3 µg ml^-1^) was added to the respective media to study the effect of the same on antimicrobial susceptibility, it ^23^. The experiments were repeated six times for consistency.

### Intracellular antibiotic accumulation assay

The accumulation assays for Norfloxacin, Ofloxacin and Bocillin FL were conducted based on the protocol previously described with some specific modifications ^41,57^. The cells expressing *rv1250* were grown till OD_600_∼0.6 and subsequently washed thrice with 50mM potassium phosphate buffer (PPB). The *E. coli* cells were energised for 20 minutes at 30°C with 0.2% glucose. The experimental and control cells were treated with 10 mg l^-1^ of fluoroquinolones and 10 µM of Bocillin FL, respectively. After every time interval (5 min), aliquots of the samples (0.5 ml) were withdrawn and washed thrice with PBB. To check for the active efflux and energy source in the efflux process, CCCP (3 µg ml^-1^) was added to the cell suspensions after 15-18 min, and aliquots were drawn further. The samples were resuspended in 0.1 M glycine hydrochloride buffer (pH 3.0) and incubated at 37°C for 2 hours for *E. coli* and overnight for *M. smegmatis* cells. The cells were further centrifuged and the fluorescence of the supernatant was assessed by using a spectrofluorimeter (FluoroMax 4, HORIBA Scientific Instruments, Irvine CA, USA) for norfloxacin (at excitation wavelength 281 nm and emission wavelength 447 nm), ofloxacin (excitation wavelength of 280 nm and emission wavelength of 485 nm) and for Bocillin FL (excitation and emission wavelength for was 488nm and 530nm, respectively). The Relative efflux (RE) was determined with the control strain using the formula: RE 1+ (N_control_ – N_test_)/N_control_. N_control_ represents the antibiotic accumulated by the control strain harbouring solely the vector control, and N_test_ represents the antibiotic uptake by the cells harbouring Rv1250 ^41,57^.

### EtBr efflux assay

The overnight cultured samples (100 µl) were washed thrice and inoculated in a 100 ml 7H9 medium. The samples were incubated at 37°C with a shaking till the culture reached an OD_600_ of ∼0.2. The *M. smegmatis* cultures were induced with 20 ng ml^-1^ of tetracycline to facilitate gene expression and protein synthesis, and incubated for 24 h, and subsequently harvested by centrifuging at 3100 g for 20 min at room temperature. The pellets obtained were washed twice with 50 mM of PPB supplemented with 1 mM MgCl_2_ and resuspended in PPB with an adjusted final OD_600_ ∼ 1.0. After keeping the cells at rest for 30 min, 2 ml aliquots of the samples were mixed with 3 µg ml^-1^ CCCP and allowed resting further for 30 min. EtBr (1.5 µM) was added to the samples and incubated at 37°C with a shaking for 4 hr. The cells were spun down at 3100 g for 20 min after keeping them at rest for 30 min, and the supernatant was discarded and resuspended at PPB. The cells were diluted by 10-fold and the fluorescence of the suspension was promptly monitored for 50 s (at an excitation wavelength of 530 nm and an emission wavelength of 600 nm) in a spectrofluorimeter (FluoroMax 4, Horiba Scientific, Japan). EtBr efflux was initiated by fast energisation using 0.4% glucose and observed for an additional 200 s. The assay was performed in triplicate and repeated thrice ^42^.

### Site-directed mutagenesis

Rv1250 cloned in pBAD-18 Cam vector was used as a template for site-directed mutagenesis. Three amino acid substitutions, Q409A, Q409E and D34A, were performed utilising specific primers (**Table S2**) and *Pfu* Turbo Polymerase (Agilent Technologies, Santa Clara, CA, USA). The amplicons were then subsequently treated with *Dpn*I (New England BioLabs, Ipswich, Massachusetts, United States) and transformed with *E. coli* XL1-Blue cells. The mutations were validated by sequencing performed by Eurofins Scientific (Hyderabad, TS, India). Further, the mutated Rv1250 was cloned in the pMIND vector and sequenced for confirmation. Confirmed mutant constructs were electroporated into *M. smegmatis* MC^2^155 cells for expression to carry out the experiments for validating the effect of mutations in the host cells.

### Semi-quantitative biofilm formation assay

*M. smegmatis* and *E. coli* cells were grown in 7H9 and LB medium, respectively. The cultures were washed with Phosphate Buffer Saline (PBS) thrice and resuspended in the aforesaid medium. Each well of a 24-well plate filled with 1 ml of 1XM63 and 1/5^th^ LB medium was inoculated with *M. smegmatis* and *E. coli* cultures (OD_600_ ∼0.6), respectively. The plates were incubated at 37°C for 48 hr. (*E. coli*) and 3 days (*M. smegmatis*). Further, the planktonic cells were removed, and the biofilm was washed thrice with PBS and kept for drying. Crystal violet (CV) solution (0.1% w/v) was added to each well of the plate and incubated for 15 min. The CV stain was removed, and the wells were washed with double-distilled water. The CV-stained wells were destained with 1 ml of 33% glacial acetic acid. The absorbance of the destained solution was measured using a micro-plate reader (Biorad iMark™ Microplate Absorbance Reader, United States) at 600 nm. Biofilm formation index was calculated by: BFI: (OD_600_ of acetic acid solubilized CV of culture – OD_600_ of acetic acid solubilized CV of control) / OD_600_ of planktonic cells. To determine the effect of CCCP on the biofilm-forming ability of the cells, the experiment was repeated in the presence of a sub-inhibitory concentration of CCCP. Each well was filled with medium supplemented with a sub-inhibitory concentration of CCCP and inoculated with cells expressing *rv1250* and control cells. The plates were incubated for 48 hr. and 3 days for *E. coli* and *M. smegmatis,* respectively. Following incubation, the wells were washed and stained with 0.1% CV and destained with 33% glacial acetic acid. The absorbance at 600nm of the destained solution was noted, and the BFI was calculated ^53,58,59^. To determine the relative carbohydrate and protein content in the extracellular matrices of biofilms, *M. smegmatis* cells of OD_600_∼ 0.6 were inoculated in a 10 ml M63 broth and incubated at 37°C in an incubator-shaker with a shaking speed of 100 rpm for 5 days. Dense biofilm layers on the walls of the tube were noted. Further, the biofilm was scraped from the walls and inoculated in a fresh M63 medium and incubated at 37°C with shaking as before for 3 days. Thereafter, the cell free broth was removed, and the biofilm layers formed was washed with PBS to remove excess salts and sugar. The biofilm layer from the walls of the tube was dissolved in Chloroform: Methanol mixture (2:1) and the mixture was centrifuged at 5000 g for 10 min. The mixture separates into a hydrophilic and hydrophobic phase. The phases were separated and used for protein and carbohydrate quantification ^60^. The carbohydrate content was estimated using the phenol-sulphuric acid assay method ^61^. Briefly, 20 µl of the samples were mixed with 20 µl of 5% phenol and 200 µl of H_2_SO_4_. The reaction mix was incubated in a dancing shaker for 5 min. The colour change of the samples was determined by obtaining absorbance at 490 nm. The carbohydrate content was determined from a standard curve of glucose. The protein concentration was estimated by the Bradford method ^62^ by mixing 50 µl of the sample with 150 µl of phosphate buffer and 2 ml of Bradford reagent. The reaction mix was incubated in the dark for 10 min. Protein content was determined by obtaining the absorbance at 620 nm and plotting the values against a BSA standard curve.

### Sur**v**ival of mycobacteria inside the macrophage cells

RAW 264.7 murine macrophage cell line was cultured and maintained in 5% CO_2_ at 37°C in Dulbecco’s modified Eagle medium (DMEM; HiMedia) enriched with 10% fetal bovine serum (FBS; Gibco, Invitrogen Life Technologies) and 1% penicillin-streptomycin solution (Sigma-Aldrich). The RAW cells were seeded in 24-well plate 24 hours prior to the infection at the density of 2.5 x 10^5^ cells per well. The media was replaced with fresh DMEM without penicillin-streptomycin. Actively growing mycobacterial cells were washed with PBS and used for infecting the macrophage cells at an approximate multiplicity of infection (MOI) of 10:1 in the presence and absence of sub-inhibitory concentrations of rifampicin (0.1 mg l^-1^) and CCCP (3 µg ml^-1^). The plates were incubated at 37°C in 5% CO_2_ for a period of 4 hr. After incubation, the wells were washed twice with PBS to remove the non-infecting bacteria. Fresh medium was added to each of the wells, and the plates were incubated further for 24 hr. After washing the cells with PBS twice, the RAW cells were lysed with 0.1% Triton X-100. The lysates were serially diluted, plated onto LB agar plates and incubated for 3 days at 37°C. The viable bacteria were counted and expressed in form of Colony Forming Unit per milliliter and data is presented as a relative survival of mycobacteria with respect to cells expressing Rv1250 ^49,53^.

### *Insilico* analysis

The model structure of the Rv1250 was retrieved from the AlphaFold database, and the strcturrs of the mutants were prepared using MODELLER. The docking was performed with Autodock Vina. All molecular dynamic simulations were performed in GROMACS 2024.1, available in the PARAM-SHAKTI supercomputer of IIT Kharagpur ^63–68^.

### Statistical Analysis

All experiments were performed in triplicate, and the results were calculated as mean±standard deviation. The statistical significance (P value) was calculated using GraphPad by performing a sample, unpaired Student’s *t*-test where *, P<0.06 (non-significant);**, P<0.006 (significant), ***P<0.0006 (significant).

## Supporting information

Rv1250_Supplementary Material_BioRxiv-Final

## Acknowledgement

We thank Dr Dasarathi Das, Scientist F, ICMR - Regional Medical Research Center, Bhubaneswar, India, for the genomic DNA of *Mycobacterium tuberculosis* H37Rv. We also thank the supercomputing facility (PARAM SHAKTI) of IIT Kharagpur.

## Author’s contribution statement

DC was involved in designing and conducting experiments, and writing the draft of the manuscript, APP was involved in *insilico* work, ARS was involved in infection studies, phylogenetic tree preparation and data analysis, SR was involved in infection studies, SBDG was involved in infection studies, conceptualiztion, draft preparation and ASG was involved in the overall supervision of the work, conceptualization, designing and editing of the manuscript.

