## Supplementary material for "Putative MFS transporter Rv1250 of *Mycobacterium tuberculosis* is involved in multidrug efflux activity": Rv1250_Supplementary Material_BioRxiv-Final

**Title:**

**Figure S1: Expression of putative MFS transporter Rv1250 in *E. coli*:**

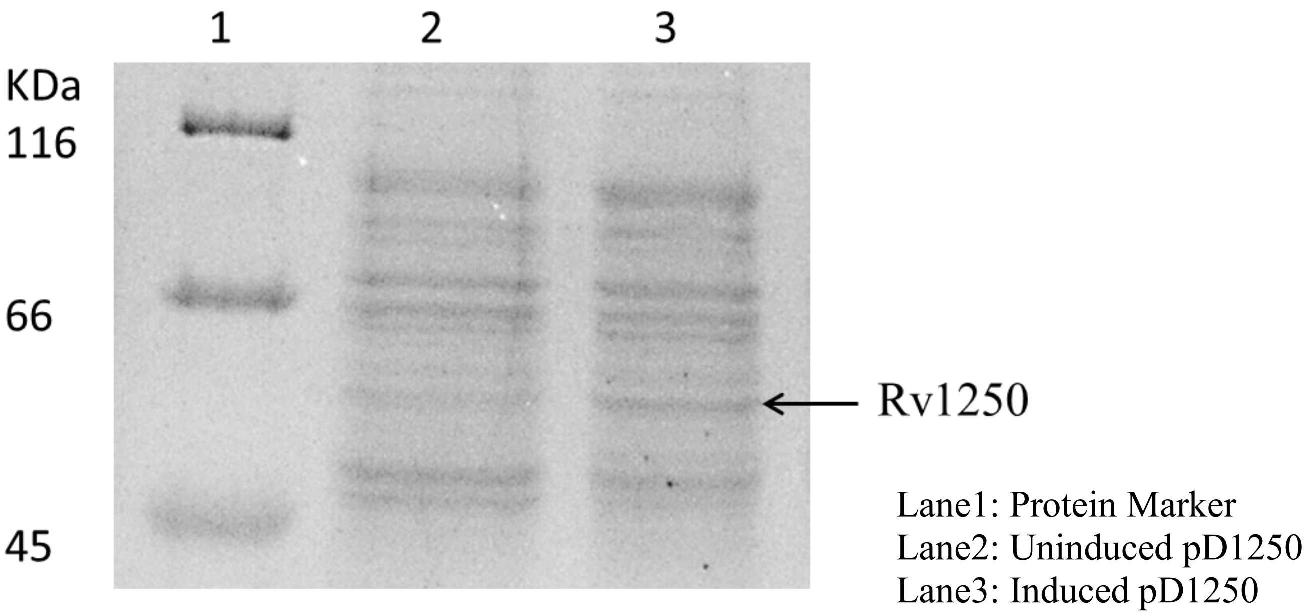


Rv1250-pBAD18Cam (pD1250) was induced and overexpressed with 0.2% arabinose in *E. coli* CS1O9 cells.

**Figure S2: Growth curve analysis:**

Growth curve analysis of *M. smegmatis* cells harbouring Rv1250, its mutants and empty pMIND vector was done following the protocol as discussed previously (1). Briefly, 7H9 medium supplemented with 0.05% Tween 80 and 100 µg/ml hygromycin was inoculated with $OD_{600}$ ~0.04 of the overnight-grown control and experimental *M. smegmatis* cells. The effect of the presence of a sub-inhibitory concentration of norfloxacin (0.25 µg $ml^{-1}$) on the cells was also measure using was determined by measuring the optical density at 600 nm using a Microplate reader (Biorad iMark™ Microplate Absorbance Reader, United States).

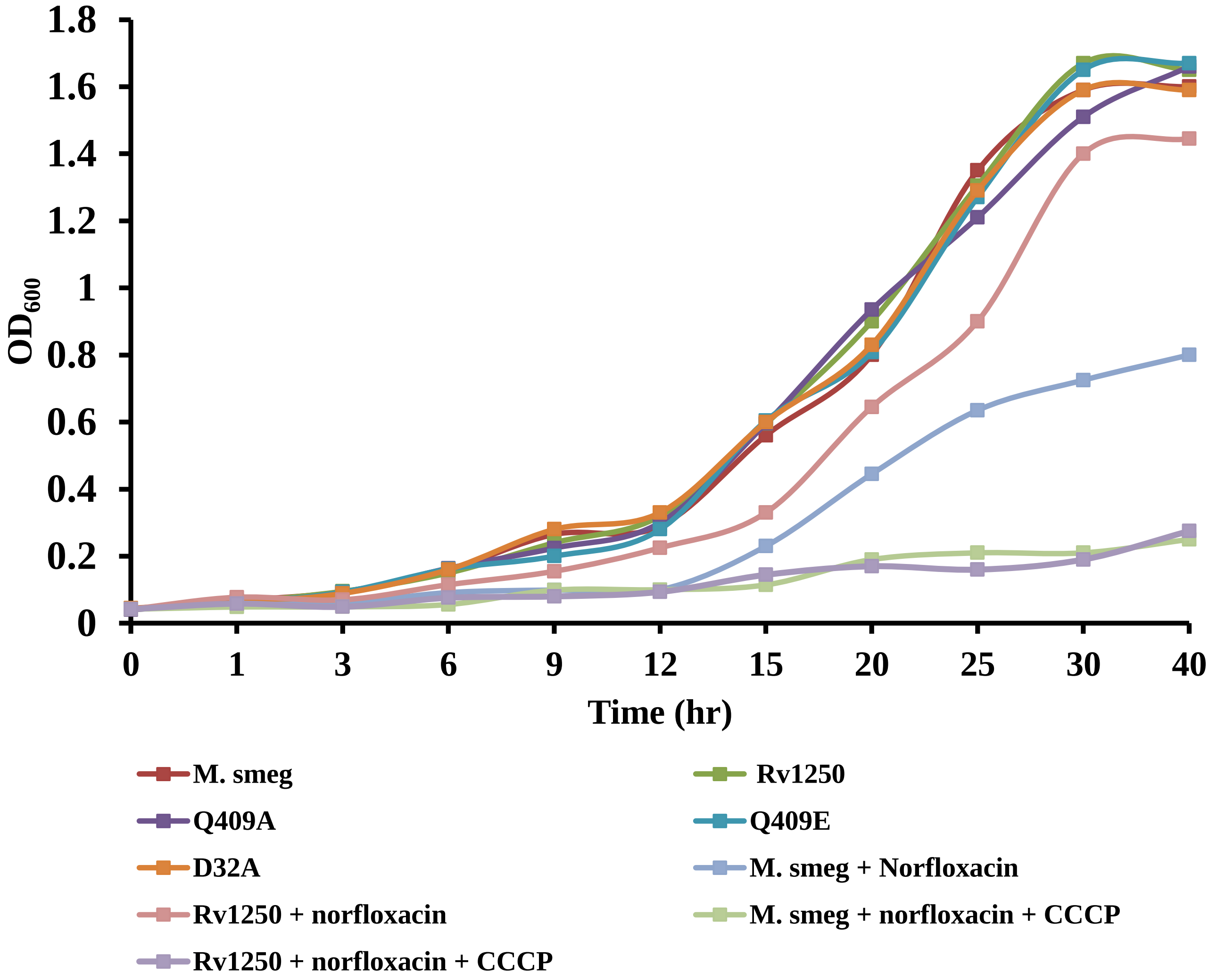


Growth curve analysis of 20 ng $ml^{-1}$ tetracycline induced *M. smegmatis* cells expressing Rv1250 in the presence of 3 µg $ml^{-1}$ and 0.25µg $ml^{-1}$ of norfloxacin.

**Table S1:** Minimum Inhibitory Concentrations of various unrelated classes of antibiotics against CS109 cells carrying pBAD18 Cam vector (control of *E. coli* cells), and CS109 cells harbouring Rv1250 (pD1250) in the presence and absence of CCCP

.

| Antibiotics | Minimum Inhibitory Concentration (MIC) mg $l^{-1}$ | | | |
|---|---|---|---|---|
| | | | CCCP | |
| | *E. coli* CS109 | CS1O9/pD1250 | *E. coli* CS1O9 | CS1O9/pD1250 |
| Norfloxacin | 0.062 | 0.25 | 0.031 | 0.062 |
| Ofloxacin | 0.062 | 0.25 | 0.031 | 0.031 |
| Levofloxacin | 0.05 | 0.1 | 0.025 | 0.05 |
| Ampicillin | 16 | 64 | 8 | 8 |
| Oxacillin | 8 | 32 | 4 | 4 |
| Amikacin | 4 | 8 | 2 | 2 |
| Apramycin | 8 | 16 | 4 | 4 |
| Gentamicin | 1 | 4 | 0.5 | 1 |

*CCCP: Carbonyl Cyanide m-Chloro-Phenylhydrazone

**Figure S3: Intracellular accumulation of norfloxacin in *E. coli:***

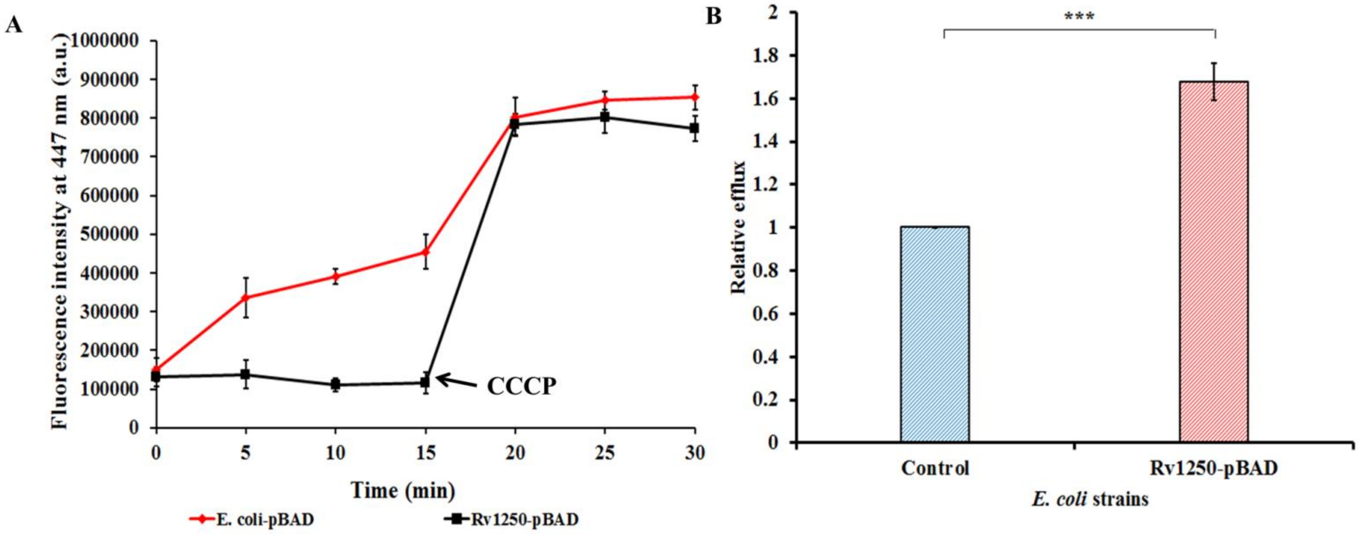


Accumulation of Norfloxacin by the cells of *E. coli* CS109 (A) harbouring Rv1250 in comparison to control cells (harbouring empty vector) with respect to time. Norfloxacin was added at $0^{th}$ min. Sample aliquots were drawn at every 5 min intervals for 30 min, and CCCP was added at $15^{th}$ min. (B) Relative efflux was calculated after exposing the *E. coli* cells to norfloxacin for 15 min. Two-tailed unpaired Student's *t*-test was performed with the control and test datasets, where ***denotes $P<0.0006$. The error bars indicate mean±SD.

**Figure S4: Phase-contrast microscopic analysis of biofilm formation of *M. smegmatis* cells:**

The 24-well plate polystyrene plate wells, consisting of cover glass slips, were filled with 1XM63 medium inoculated with $10^5$ cells and incubated at 37°C for 3 days under static conditions. The wells were washed with PBS buffer and stained with 0.1% crystal violet solution for 15 min, and finally, the wells were washed with water and dried. The biofilms formed on the coverslips were subsequently visualised using an OLYMPUS 1×73 (2)

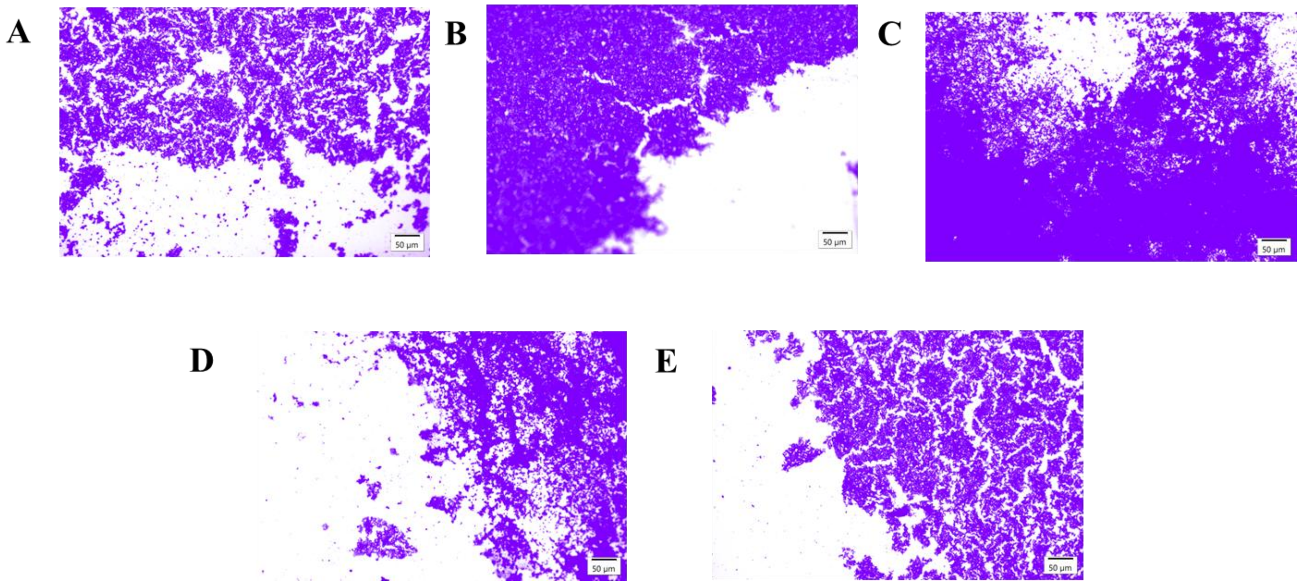


**Fig A:** CV stained wild-type *M. smegmatis* biofilm **Fig B:** CV stained pM1250 **Fig C:** CV stained pM1250-Q409A **Fig D**: CV stained pM1250-Q409E **Fig E**: CV stained pM1250-D32A

Microscopic analysis of CV-stained biofilms: Bright-field microscopic images (10×) of the biofilms formed at the air-liquid interface after staining with CV. The scale bar represents 50 µm.

**Figure S5: *In silico* analysis of Rv1250 and its mutants**

### Structure modelling and MD simulation:

The structural model of Rv1250 was sourced from the AlphaFold database and subsequently modified by eliminating 16 amino acids from the N-terminal signal peptide and 96 from the C-terminal region (3). All the structures were energy-minimised using 1,000 steps of steepest descent and 100 steps of the conjugate gradient method, employing the AMBER-ff14SB force field in Chimera software (4).

Ligand docking was performed using AutoDock Vina, and the systems of the native protein and three mutated protein-ligand complexes (D34A, Q409A and Q409E) were prepared from the native docking complex using CHARMM_GUI for molecular dynamics simulation (5, 6). The protein-ligand complex was embedded in a POPE membrane bilayer and parameterised with CHARMM36 and CGenFF force fields (6). It was solvated with TIP3 water molecules and neutralised $Na^+$ and/or $Cl^-$ ions. Energy minimisation was conducted in GROMACS 2024.1 until the maximum force reached 1,000 $kJ/mol/nm^{-2}$ (7). Equilibration was performed using NVT and NPT ensembles for 1 ns at 300 K, with the protein backbone position-restrained. Production simulations followed for 500 ns with a 2-fs time step, saving coordinates every 250 ps. The V-rescale thermostat kept the temperature at 300 K, while the Parrinello-Rahman barostat maintained pressure at 1 bar. LINCS was used for hydrogen bond constraints, and long-range electrostatic interactions were calculated via the Particle Mesh Ewald method with a 1.2 nm cutoff for Coulombic and van der Waals interactions.

Two enhanced sampling methods were employed separately: umbrella sampling (US) and the extended Adiabatic Biasing Force (eABF) method, implemented using the COLVAR module in GROMACS 2024.1 (8). The collective variable for enhanced sampling was defined as the Z-axis distance between the centre of gravity of the protein (aligned along the channel axis) and the docked ligand molecule. For umbrella sampling, 40 equally space windows were used, and the WHAM program was applied to calculate the potential of mean force (PMF). For the eABF simulation, the “full sample”, “extended fluctuation”, and “width” parameters were set to 4000, 0.1 and 0.5, respectively.

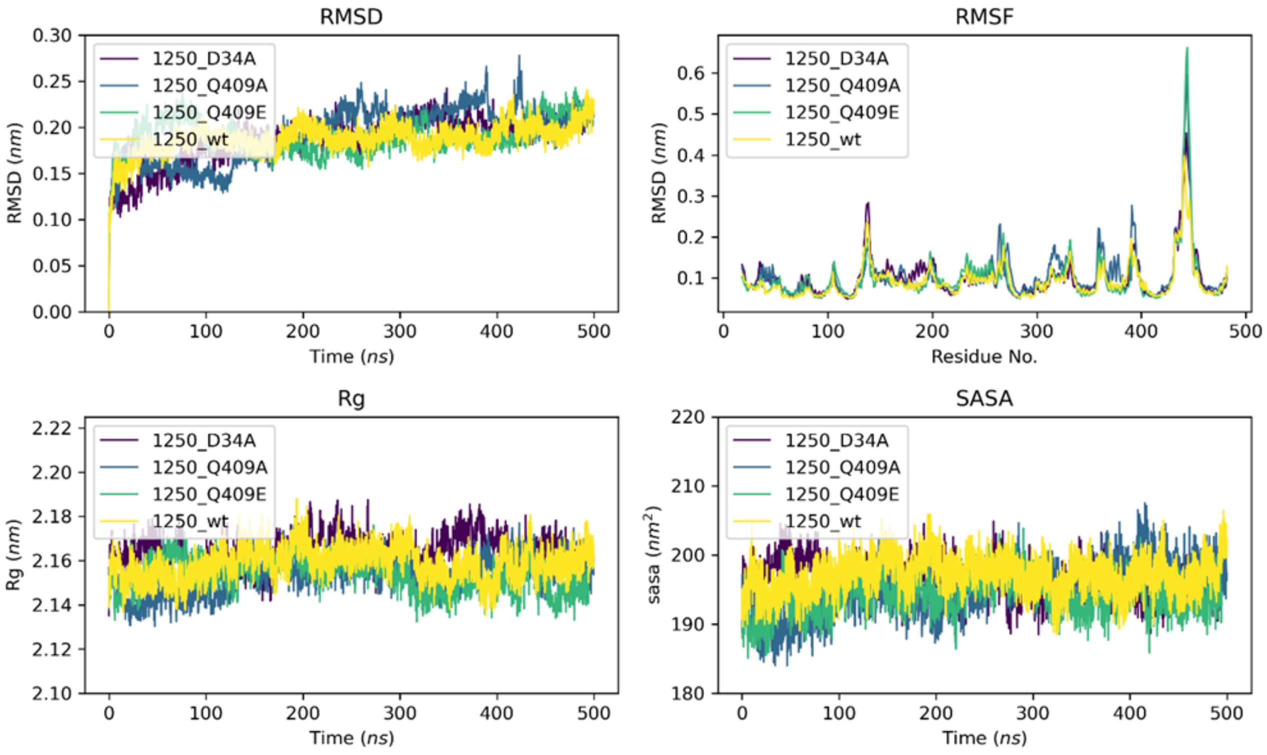


Comparison of RMSD, RMSF, Rg and SASA for wild-type and mutant protein (D34A, Q409A and Q409E).

**Table S2:**

| **Primers** | **Sequences (5'-3')** |
|---|---|
| **Rv1250/pMIND/FP** | CTCTCTCATATGAGGAGGCTCTCTCTATGACTACGGCGATAC |
| **Rv1250/pMIND/RP** | CTCTCTACTAGTTTATAGTGCGCTCGGAGCTGTTGACTCAGTATC |
| **Rv1250/pBAD18Cam/FP** | CTCTCTGCTAGCAGGAGGCTCTCTCTATGACTACGGCGATAC |
| **Rv1250/pBAD18Cam/RP** | CTCTCTAAGCTTTTATAGTGCGCTCGGAGCTGTTGACTCAGTATC |
| **Rv1250/Q409E/FP** | TTCAACGCCGTCCGGGAGCTGGGGGCTGTGCTG |
| **Rv1250/Q409E/RP** | CAGCACAGCCCCCAGCTCCCGGACGGCGTTGAA |
| **Rv1250/Q409A/FP** | TTCAACGCCGTCCGGGCGCTGGGGGCTGTGCTG |
| **Rv1250/Q409A/RP** | CAGCACAGCCCCCAGCGCCCGGACGGCGTTGAA |
| **Rv1250/D34A/FP** | GGCGGCGCGCTGGTCGCCAGCATGGGGTGGGAG |
| **Rv1250/D34A/RP** | CTCCCACCCCATGCTGGCGACCAGCGCGCCGCC |
